# The vaccine candidate Liver Stage Antigen 3 is exported during *Plasmodium falciparum* infection and required for liver-stage development

**DOI:** 10.1101/2025.07.18.665593

**Authors:** Robyn McConville, Ryan W.J. Steel, Matthew T. O’Neill, Alan F. Cowman, Norman Kneteman, Justin A. Boddey

## Abstract

*Plasmodium falciparum* remodels infected erythrocytes using exporting effector proteins. Parasites express the aspartyl protease plasmepsin V that processes proteins containing a PEXEL motif and the PTEX translocon to successfully export proteins. During liver-stage infection, PTEX is required for *P. falciparum* development, but which proteins are exported remain largely unclear, yet they may serve important functions and be presented by MHC-I molecules, thereby representing potential vaccine candidates. Here, we investigated liver stage antigen 3 (LSA3), an immunogenic protein of the Laverania subgenus of *Plasmodium*. We show that LSA3 possesses a PEXEL motif processed by plasmepsin V and is targeted to one or more membranes surrounding the blood-stage parasite, suggestive of the parasitophorous vacuole membrane (PVM). A subset of LSA3 also localizes in the erythrocyte, where it forms punctate structures that are not Maurer’s clefts but are soluble in biochemical fractionation assays reminiscent of J-dot proteins. During infection of human hepatocytes, antibodies to LSA3 co-localize with EXP1 and EXP2 at the PVM yet these antibodies were not detected beyond this membrane. Finally, genetic disruption of *LSA3* in *P. falciparum* NF54 attenuated fitness at the liver stage, manifesting as a 40% reduction in parasite liver load by day 5 postinfection of humanized mice. The identification of LSA3 as a member of the *P. falciparum* exportome and important for liver-stage development confirms the hypothesized potential of exported proteins as promising vaccine candidates, underscoring the need for their continued discovery and biological characterization, including those expressed at the liver stage.

## Introduction

*Plasmodium* species are protozoan protists responsible for more than 282 million cases of malaria and 610,000 deaths each year (WHO, 2025). While major progress has been made in reducing the incidence of malaria over the last decade, these advances are threatened by development of resistance to front-line antimalarials. The development of next generation therapies and vaccines require a better understanding of how these parasites survive and proliferate in the vertebrate and mosquito hosts (Kappe *et al*, 2010). This includes mechanisms for inhabiting different host cell types, evasion of immune responses and their ability to acquire nutrients to sustain the extraordinary growth rates in the liver for onward transmission (Vaughan & Kappe, 2017). The liver-stage is currently the target of the only approved vaccines, though they offer modest efficacy (Clinical Trials Partnership, 2015; Datoo *et al*, 2021), highlighting the need for continued research.

Infections are initiated by *Plasmodium* sporozoites injected into the skin by an infected mosquito, which migrate to the liver and develop asymptomatically in hepatocytes for ∼7-10 days (Frischknecht & Matuschewski, 2017). Liver merozoites subsequently egress the liver and initiate cycles of infection in red blood cells (RBC) that are responsible for clinical malaria. Both the liver and RBC stages of *Plasmodium* replicate within a unique parasitophorous vacuole membrane (PVM) and have evolved mechanisms to remodel this membrane and their host cells (Kappe *et al*., 2010). During the blood stage, this is achieved by the export of proteins to or across the PVM into the infected erythrocyte. For many exported proteins, their targeting is dependent on the *Plasmodium* export element (PEXEL) / host targeting (HT) motif (RxLxE/Q/D) (Hiller *et al*, 2004; Marti *et al*, 2004) that is recognized by the endoplasmic reticulum (ER)-resident aspartyl protease plasmepsin V (Boddey *et al*, 2010; Russo *et al*, 2010). This protease processes the PEXEL between the third and fourth residues (Boddey *et al*, 2009; Chang *et al*, 2008), licensing the matured and then N-acetylated proteins for export. A subset of PEXEL-negative proteins (PNEPs) is also exported into RBCs (Heiber *et al*, 2013; Spielmann *et al*, 2006). Export across the PVM is mediated by the *Plasmodium* translocon of exported proteins (PTEX) (Beck *et al*, 2014; de Koning-Ward *et al*, 2009; Elsworth *et al*, 2014; Ho *et al*, 2018). PEXEL cleavage by plasmepsin V can also target proteins to alternate locations such as the parasitophorous vacuole and rhoptries (Fierro *et al*, 2024; Freville *et al*, 2024).

The nucleated hepatocyte presents a vastly different cellular environment, including a complex secretory network and cell intrinsic immune programs including interferons, guanylate binding proteins (GBPs), lysosomes and autophagic proteins that can target the intracellular parasite niche; hepatocytes also use MHC presentation pathways for recruiting adaptive immune responses that can ultimately attack and clear the parasite (Cockburn *et al*, 2011; Liehl *et al*, 2014; Marques-da-Silva *et al*, 2022; Marques-da-Silva *et al*, 2025; Prado *et al*, 2015; Weiss, 1990). Whether or to what extent parasite liver-stages translocate their own proteins into or beyond the PVM to diminish hepatocyte defences remain largely unanswered. However, several lines of evidence suggest that a protein export pathway involving components characterized in blood-stages is expressed and is functionally important in liver-stage parasites, but with some important differences.

Firstly, the PTEX150 and EXP2 components of the PTEX translocon are expressed at the parasite periphery of, and are important for, *P. falciparum* growth in hepatocytes, suggesting export of proteins may occur (McConville *et al*, 2024). EXP2 serves dual roles in the PVM of RBCs, both as a protein-conducting gateway for export and as a nutrient-transporter for parasite growth (Garten *et al*, 2018); its importance during hepatocyte infection may be, by analogy, for either function. The PTEX unfoldase HSP101 is expressed in *P. falciparum* liver-stages (Zanghi *et al*, 2025) but its importance remains unknown. HSP101 is apparently absent from the liver-stage translocon in the rodent malaria parasite *P. berghei* (Kreutzfeld *et al*, 2021; Matz *et al*, 2015).

Secondly, multiple PEXEL and PNEP proteins are expressed by *P. falciparum* liver-stages (Zanghi *et al*., 2025), and in *P. berghei* LISP2 is efficiently exported beyond the PVM into the infected hepatocyte cytoplasm and nucleus (Orito *et al*, 2013); interestingly, targeting of LISP2 was limited to the PVM of *P. falciparum*-infected hepatocytes *(McConville et al., 2024)* yet this protein is important for late liver-stage development of this species (Zanghi *et al*., 2025). Proteomics analysis of *P. berghei* merosomes also revealed the presence of peptides arising from PEXEL cleavage by plasmepsin V including LISP2, suggesting this protease is functional in liver-stages. Finally, while only few blood-stage PEXEL proteins are trafficked to the parasitophorous vacuole (PV), the *P. falciparum* PEXEL proteins LISP2, circumsporozoite protein (CSP) and LSA3 are all trafficked to the periphery of the liver-stage parasite during hepatocyte infection (Daubersies *et al*, 2000; McConville *et al*., 2024; Zanghi *et al*., 2025) and genetic disruption of PTEX subunits reduced expression of LISP2 and CSP, suggesting a direct or indirect link between the translocon and protein trafficking and/or quality control (McConville *et al*., 2024).

Only the Laverania subgenus of *Plasmodium* possess the *LSA3* gene; in the most virulent human parasite, *P. falciparum*, LSA3 is expressed during infection of hepatocytes (Daubersies *et al*., 2000) and RBCs (Morita *et al*, 2017). LSA3 contains an N-terminal putative PEXEL motif (Maier *et al*, 2008; McConville *et al*., 2024; Morita *et al*., 2017), suggesting it may be an exported protein, yet whether the PEXEL is functional has not been definitively proven. LSA3 also possesses an N-terminal signal anchor (that would be removed following plasmepsin V cleavage of the PEXEL motif) and a putative C-terminal hydrophobic region. LSA3 is a protective antigen in immunization studies; however, despite its immunogenicity, the amino acid sequence is highly conserved across diverse *P. falciparum* geographical isolates. This is suggestive of functional importance and strong evolutionary pressure against sequence variation (Perlaza *et al*, 2001; Perlaza *et al*, 2008), although, functional characterization of *LSA3* deletion mutants have not been reported at the mosquito or liver stages of the lifecycle and genetic disruption at the blood-stage did not identify an essential phenotype (Maier *et al*., 2008). Research efforts have instead largely focused on the immunogenicity of LSA3, with its initial discovery as a pre-erythrocytic antigen involved in protection against future liver-stage infection (Daubersies *et al*., 2000).

More recently, LSA3 was identified as a novel blood-stage antigen following a screen of a wheat germ expression library of *P. falciparum* proteins with human sera from individuals living in endemic areas, with the human antibodies that bound to LSA3 reducing erythrocyte invasion by ∼24% (Morita *et al*., 2017). In schizonts, LSA3 localizes to dense granules (DGs) (Morita *et al*, 2025; Morita *et al*., 2017) yet it is not known where it localizes earlier during erythrocytic infection. Recently, we genetically disrupted of *LSA3* in *P. falciparum* NF54 and characterization of this mutant showed it supports a role during completion of erythrocyte invasion (Morita *et al*., 2025).

LSA3 was historically characterised as a promising vaccine candidate due its protective effect against *P. falciparum* sporozoite challenge in both chimpanzee and Aotus monkey models (Daubersies *et al*., 2000; Perlaza *et al*., 2008). Immunofluorescence assays (IFA) of *P. falciparum* sporozoites using affinity-purified LSA3 human antibodies from protected volunteers given radiation-attenuated sporozoites (RAS) detected labelling at the sporozoite surface (Daubersies *et al*., 2000). IFA of *P. falciparum* 5-day old liver parasites in monkeys using antibodies induced by LSA3 lipopeptide injection of a chimpanzee detected labelling at the periphery of exoerythrocytic parasites, altogether suggesting LSA3 localizes at the PVM (Daubersies *et al*., 2000). However, confirming this location is challenging and requires specific PVM markers that have not been used to date, nor have LSA3-deficient parasites been characterized at the liver stage.

Here, we have undertaken a biochemical and functional study of LSA3 in *P. falciparum* across the blood, mosquito and liver stages. Using LSA3-C and LSA3-T antibodies (Morita *et al*., 2025; Morita *et al*., 2017) epitope tagging, and inhibitors, we demonstrate that LSA3 has a *bona fide* PEXEL sequence that is proteolytically processed by plasmepsin V and that LSA3 is targeted both to the PVM and into the infected erythrocyte cytoplasm where it forms punctate structures and is therefore confirmed to be a member of the malarial exportome (Sargeant *et al*, 2006). In humanized mice, we showed that antibodies to LSA3 co-localized with the PVM markers EXP1 and EXP2. However, we did not detect LSA3 beyond the PVM of infected hepatocytes, suggesting LSA3 is secreted to the PVM or may be considered exported if a domain is exposed to the hepatocyte lumen from this membrane. Genetic disruption of *LSA3* in *P. falciparum* NF54 did not prevent gametocytogenesis, transmission to or development within the mosquito; however, LSA3-deficiency caused a significant reduction of parasite fitness by day 5 of the liver stage in humanized mice, demonstrating that LSA3 is important for *P. falciparum* liver stages.

## Results

### LSA3 is exported to the *P. falciparum*-infected erythrocyte

LSA3, encoded by <u>PF3D7_0220000</u>, is conserved in all primate-infective *Plasmodium* species and contains an N-terminal hydrophobic signal anchor likely for ER entry, a PEXEL motif followed by disordered or flexible repeat regions and a predicted C-terminal hydrophobic region. AlphaFold predicts a domain (residues 908-1115) with some similarity to the substrate binding domain of the chaperone, DnaK (Daubersies *et al*., 2000; Jumper *et al*, 2021; Morita *et al*., 2017; Prieur & Druilhe, 2009) (***Figure 1A***). The presence of an N-terminal hydrophobic sequence and PEXEL suggested LSA3 may be a substrate for the ER-resident aspartyl protease, plasmepsin V, raising the possibility it is exported (Marti *et al*., 2004; McConville *et al*., 2024). Despite initially being discovered as a liver-stage protein (Daubersies *et al*., 2000), LSA3 is also expressed in blood stage merozoites where it localises to DGs and is required for completion of RBC invasion, but its localization in ring and trophozoite stages has not yet been characterized (Maier *et al*., 2008; Morita *et al*., 2025; Morita *et al*., 2017).

**Figure 1.**
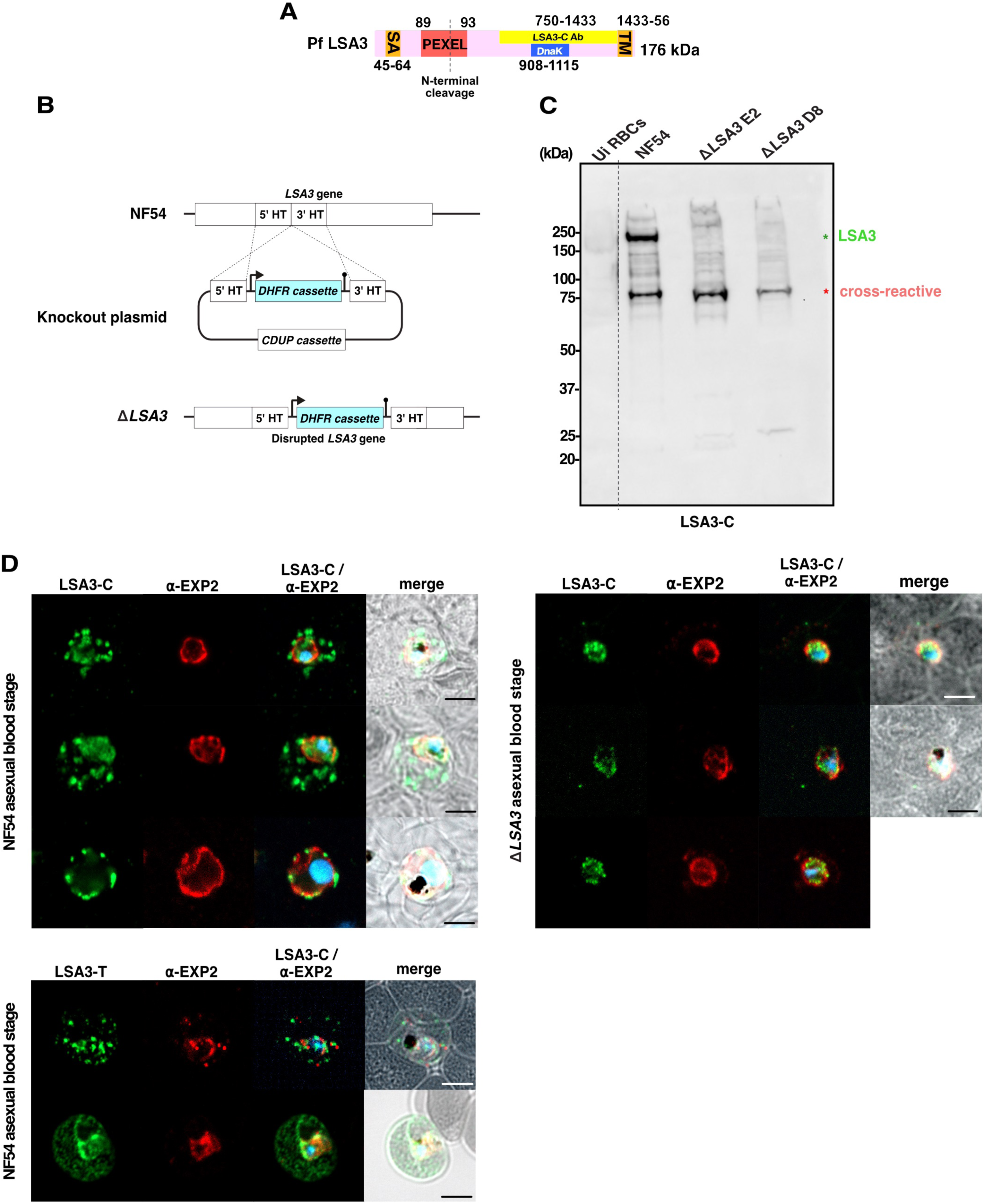
Genetic disruption of *LSA3* in *P. falciparum*. **A.** Schematic of the 176 kDa LSA3 protein conserved across human-infective malaria species. Regions between residues 45-64 and 1433-1456 are predicted to be a hydrophobic signal anchor (SA) and transmembrane domain (TM) (orange), respectively; residues 89-93 contain a PEXEL motif (red) and residues 908-1154 have homology to the substrate binding site from the chaperone DnaK (blue). LSA3-C antibody (Ab) binds the C-terminal region (yellow). **B.** Schematic of *LSA3* genetic disruption strategy in *P. falciparum* NF54. The endogenous coding sequence of LSA3 was disrupted by allelic exchange with a selection cassette encoding a 5’ promoter-dihydrofolate reductase gene (*DHFR*)-3’ terminator flanked by homology targets (HT) internal to the *LSA3* gene, causing disruption of protein expression. Negative selection of the remaining knockout construct was performed with a cytosine deaminase and uracil phosphoribosyltransferase (*CDUP*) cassette by addition of 5-FC to parasite cultures. **C**. Immunoblot of uninfected human red blood cells (Ui RBCs) or parasite blood-stage lysates of NF54 (parental control) and LSA3 KO clones E2 and D8, probed with LSA3-C antibodies. **D.** IFA of NF54 parental parasites following methanol/acetone fixation and staining with LSA3-C, LSA3-T or anti-EXP2 (PVM) antibodies. IFA of Δ*LSA3* parasites using LSA3-C and anti-EXP2 antibodies confirms loss of exported LSA3 in infected erythrocytes and non-specific binding to one or more proteins inside the parasites. Representative images of n=3 experiments. Scale bars, 5 µm.

Given the possibility of being exported and the strong immunogenicity of LSA3, we functionally characterised it by generating transgenic *P. falciparum* NF54 Δ*LSA3* clones, in which the *LSA3* gene was disrupted by allelic exchange (***Figure 1B***). Integration of the Δ*LSA3* knockout plasmid was confirmed in independent clones by polymerase chain reaction (PCR) and clones D8 and E2 were selected for further analysis (***Figure S1***). Loss of LSA3 protein expression was confirmed by immunoblot with antibodies recognizing the LSA3 C-terminus (LSA3-C) for both knockout clones D8 and E2 (***Figure 1C***); a non-specific band at ∼75 kDa was present, which was of parasite origin as it was absent in uninfected RBCs. While LSA3 has a predicted molecular weight of 175 kDa, it migrates at a larger size in SDS-PAGE (Morita *et al*., 2017). Validation that *LSA3* was successfully disrupted in NF54 was also provided using the recently generated LSA3-T antibody, which is not cross-reactive, in an accompanying manuscript (Morita *et al*., 2025).

To further validate loss of LSA3 expression, IFA of RBCs infected with either NF54 parental or Δ*LSA3* E2 parasites was performed using LSA3-C (***Figure 1D***). LSA3-C localized both inside the parasite and beyond the EXP2-defined PVM in punctate structures indicating it was an exported protein (***Figure 1D***). By contrast, in all erythrocytes infected with Δ*LSA3* parasites, the exported signal was absent, confirming LSA3 is exported and the specificity of LSA3-C, but internal puncta remained detectable, likely representing the cross-reactive species observed by immunoblot at ∼75 kDa (***Figure 1C, D***). IFA with the recently generated LSA3-T antibody (Morita *et al*., 2025) further confirmed that LSA3 was exported into the erythrocyte in trophozoites (***Figure 1D***).

Interestingly, while antibodies that react with LSA3 were shown to reduce merozoite invasion of erythrocytes by ∼24% in growth inhibitory assays (Morita *et al*., 2017), we were able to generate deletion mutant clones in NF54, similar to previous studies using 3D7 (Maier *et al*., 2008; Oberstaller *et al*, 2025) confirming that LSA3 is not critical here; however its genetic disruption in NF54 did slow the rate of erythrocyte invasion by merozoites and led to the presence of acollé forms, as recently reported (Morita *et al*., 2025). This demonstrates that LSA3 supports the completion of invasion by *P. falciparum* merozoites yet is not critical for blood-stage growth while, in addition to DG localization, LSA3 is expressed in trophozoites where it is exported into the infected erythrocyte and is located within punctate structures.

### LSA3 does not localize to Maurer’s cleft tethers

*P. falciparum* exports proteins into distinct punctate structures in the iRBC, including Maurer’s clefts that become tethered to the erythrocyte membrane, J-dots (Kulzer *et al*, 2010) and electron-dense vesicles (EDVs) that are mobile structures within the erythrocyte cytosol. IFA with antibodies to LSA3 and membrane associated histidine rich protein 2 (MAHRP2), which localizes at Maurer’s cleft tethers (Pachlatko *et al*, 2010) indicated that although antibodies to both LSA3 and MAHRP2 exhibit punctate signals in the RBC, no colocalization occurred, suggesting LSA3 does not localize to Maurer’s clefts or associated tethers (***Figure 2A***) but instead is located within alternate structures that were further characterized below.

**Figure 2.**
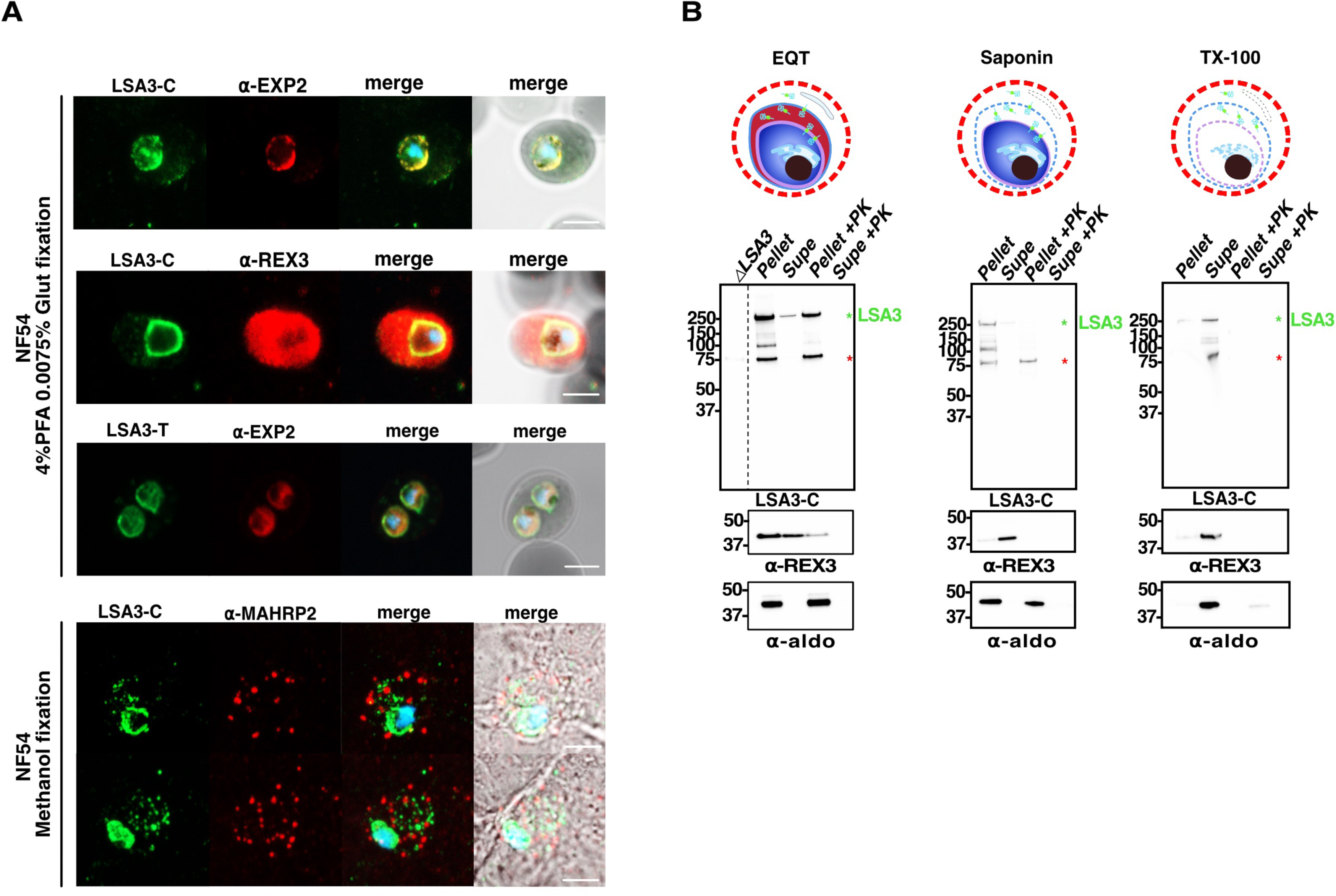
LSA3 is trafficked into the PVM and the erythrocyte during blood-stage development. **A.** IFA of NF54 parental parasites following fixation with paraformaldehyde/glutaraldehyde (bottom) and methanol/acetone (top) and staining with anti-LSA3 (LSA3-C or LSA3-T) and antibodies to EXP2 and REX3 (top) or MAHRP2 (Maurer’s cleft tethers, bottom). IFA of NF54 parasites following PFA fixation and staining with either anti-LSA3 (LSA3-C), anti-EXP2 or anti-REX3 antibodies (below). Scale bars, 5 µm. **B.** Proteinase K (PK) protection assays to localize LSA3. Erythrocytes infected with NF54 parental parasites were enriched by Percoll centrifugation and permeabilized with either equinatoxin II (EQT; selectively lyses the erythrocyte membrane), 0.03% saponin (selectively lyses the erythrocyte membrane and PVM) or 0.1% Triton-X-100 (lyses all membranes). The pellet was separated from the supernatant (supe) and both fractions were incubated with PBS containing Proteinase K (PK) at 37 °C. Protease inhibitor cocktail and PMSF was added to all samples followed by reducing sample buffer before immunoblotting with LSA3-C, anti-REX3 (exported control) or anti-aldolase (aldo; internal protein control for parasite membrane integrity) antibodies. Representative data is from n=3 experiments.

### LSA3 is membrane-associated at the PVM and soluble in the erythrocyte

To further understand the localization of LSA3, IFA with LSA3-C was performed with the exported protein REX3 as a control (Spielmann *et al*., 2006). This revealed distinct patterns of localisation, with LSA3-C forming aggregates and REX3 appearing diffuse throughout the cytoplasm as expected (Gruring *et al*, 2012; Schulze *et al*, 2015) (***Figure 2A***). Detection of REX3 required fixation with 4% paraformaldehyde (PFA) yet, under these conditions, export of LSA3 was poorly visible with LSA3-C or LSA3-T; instead, these LSA3 antibodies accumulated at the parasite periphery and co-localized with EXP2 at the PVM (***Figure 2A***). Therefore, the epitopes recognized by LSA3-C and LSA3-T were preserved with methanol:acetone fixation, which is not without precedent (Kulzer *et al*., 2010). As different fixatives were necessary, co-imaging of exported REX3 and LSA3 in the same infected erythrocyte was not possible.

To better understand the localization and solubility profile of LSA3, iRBCs were analyzed biochemically using a pore-forming toxin or detergents and protease digestion followed by Western blotting (***Figure 2B***). This approach confirmed that a soluble form of LSA3 was exported into the iRBC, as shown by release of protein into the supernatant after incubation with EQT, a pore-forming toxin from the sea anemone *Actinia equina* that selectively permeabilizes the erythrocyte membrane but not the Maurer’s clefts or PVM (Jackson *et al*, 2007). Exported LSA3 in the EQT supernatant was sensitive to PK, indicating the antibody-binding domain was exposed to the iRBC cytosol; by contrast, the EQT pellet fraction of LSA3 was mostly resistant to PK, indicating this fraction was not exposed to the RBC cytosol, consistent with a location within the parasite and/or PV/PVM (***Figure 2B***). To ensure EQT selectively lysed the erythrocyte membrane, the exported protein REX3 was assessed as a control (Spielmann *et al*., 2006). Anti-REX3 antibodies revealed that soluble REX3 was exported (liberated into the EQT supernatant) and sensitive to PK as expected (***Figure 2B***). The cytoplasmic parasite protein aldolase also remained inaccessible to EQT and PK as expected, confirming integrity of the parasite membrane (Boddey *et al*., 2009). We also confirmed that the antibody-binding domain of LSA3 was not presented on the iRBC surface, as it was resistant to PK when intact iRBC were treated with this enzyme (***Figure S2***).

Next, the subcellular localization of LSA3 within the parasite and parasitophorous vacuole was biochemically investigated using detergents. Treatment of iRBC with the plant-derived detergent saponin, which selectively permeabilizes membranes of the RBC, Maurer’s clefts and PVM, while leaving parasite membranes intact, again released exported LSA3 (the quantity released was similar to EQT alone) into the supernatant however the protein was abundant in the saponin pellet and sensitive to PK, indicating it was membrane-associated with the antibody-binding domain facing the PV lumen (***Figure 2B***). This indicated that LSA3 was predominantly located at the PVM or plasma membrane of *P. falciparum*-iRBCs. The REX3 control was predominantly in the saponin supernatant and sensitive to PK as expected, consistent with this protein being exported and within the PV prior to translocation into the host cell (***Figure 2B***). Both Aldolase and the ∼75 kDa cross-reactive band recognized by LSA3-C remained inaccessible to saponin and PK, confirming the parasite membrane was intact and corroborating the IFA analysis that this cross-reactive protein was located inside the parasite.

To better understand the membrane association of LSA3, iRBC were incubated with a low concentration of the non-ionic detergent TX-100 (Gabriela *et al*, 2022), which liberated the majority of LSA3 (and REX3 as expected) into the supernatant, with both the pellet and supe fractions now accessible to PK (***Figure 2B***). This suggests that the majority of membrane-associated LSA3 could be peripherally rather than integrally associated with membranes with the remining protein in the pellet representing a subpopulation more tightly bound to detergent-resistant structures or incomplete solubilization due to limited detergent accessibility, which was also observed for aldolase that may be bound as a complex (***Figure 2B***). Altogether these data suggest that LSA3 localizes peripherally to the luminal leaflet of the PVM, although we cannot exclude binding to the parasite membrane, and in soluble punctate structures in the infected erythrocyte.

### The PEXEL of LSA3 is processed by plasmepsin V

The observation that LSA3 is exported during the blood-stage raised the question of molecular mechanism. Previously, we and others identified a PEXEL motif in LSA3 from *P. falciparum* that is conserved in LSA3 from *P. malariae*, *P. ovali curtsi* and *P. reichenowi* (McConville *et al*., 2024; Morita *et al*., 2017; Sargeant *et al*., 2006). To test whether the PEXEL in LSA3 was functional, *LSA3* mini-gene reporters were generated that encompassed the N-terminal 112 residues of endogenous LSA3 from *P. falciparum* with a native PEXEL sequence (RSLGE) or a mutated PEXEL RLE>A (ASAGA) fused to green fluorescent protein (GFP) (***Figure 3A***). This approach was chosen because native LSA3 is a large protein of 176 kDa, potentially masking minor size shifts from N-terminal processing in immunoblots. The mini genes did not encode the C-terminal sequences recognized by LSA3 antibodies and so anti-GFP antibodies were used. The constructs were transfected into *P. falciparum* 3D7 parasites and integrated into the *p230* locus by double cross-over homologous recombination to ensure that expression of other genes required for asexual parasite growth were not disrupted and expression of the reporters was driven by the *BIP* gene promoter (***Figure 3A***) and positive clones selected via Blasticidin-S-deaminase (Ashdown *et al*, 2020).

**Figure 3.**
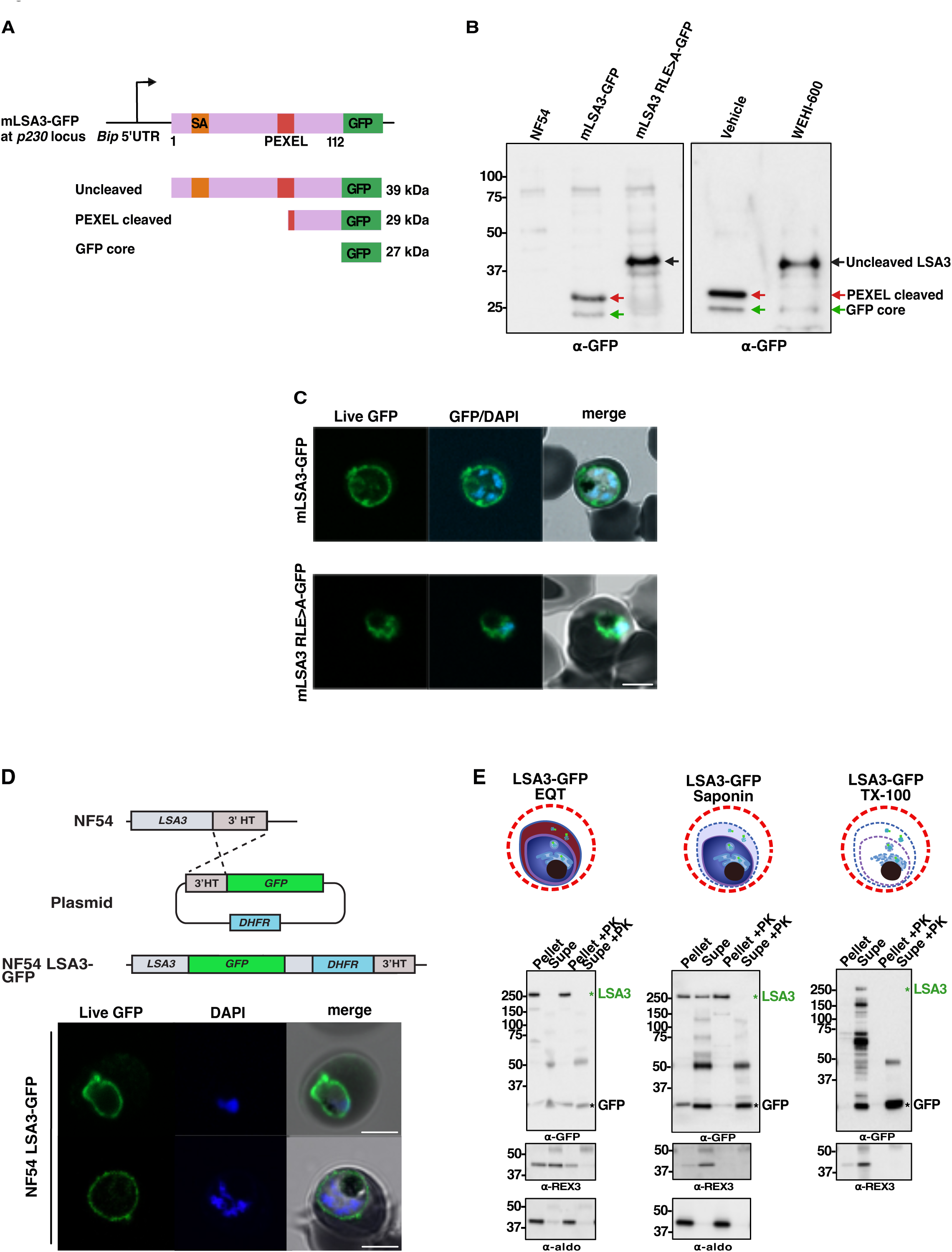
The PEXEL of LSA3 is processed by plasmepsin V. **A.** Schematic of LSA3-GFP mini gene expressed from the *BIP* gene promoter following integration into the redundant *p230* locus. Two minigenes fused to green fluorescent protein (GFP) were generated containing endogenous LSA3 residues 1-112 encoding the N-terminal signal anchor (SA) and either a native PEXEL motif (red) or with key PEXEL residues mutated to alanine (RLE>A). Predicted molecular weights following possible putative cleavage events are depicted**. B.** Immunoblots of blood-stage lysates from parental NF54 control and transgenic NF54 expressing LSA3 minigenes probed with anti-GFP antibodies. Left blow shows accumulation of uncleaved mLSA3 RLE>A-GFP compared to mLSA3-GFP and NF54 parental controls (lanes 1-3). Right blot shows accumulation of uncleaved mLSA3-GFP in the presence of plasmepsin V inhibitor WEHI-600 for 5 hr compared to DMSO vehicle control. The uncleaved band was a similar size as uncleaved mLSA3 RLE>A-GFP in the left blot. Predicted sizes: full length minigene: 39 kDa; PEXEL processing: 29 kDa; GFP core in the food vacuole following secretion of reporter: 27 kDa. **C.** Live fluorescence imaging of transgenic NF54 blood-stage parasites expressing either mLSA3-GFP (secreted to PV; top) or mLSA3 RLE>A-GFP (retained within parasite, likely the ER; bottom). Scale bars, 5 µm. **D.** Schematic of the strategy used to endogenously tag LSA3 with GFP at the C-terminus in *P. falciparum* NF54 (top). Bottom panel shows live fluorescence imaging of LSA3-GFP expressed from the endogenous locus in transgenic NF54 parasites with the protein localized at the PV and PVM (see E). Scale bars, 5 µm. **E.** Proteinase K (PK) protection assays to localize endogenous LSA3-GFP. Erythrocytes infected with NF54 LSA3-GFP were enriched by Percoll centrifugation and permeabilized with either equinatoxin II (EQT; selectively lyses the erythrocyte and Maurer’s cleft membranes), 0.03% saponin (selectively lyses the erythrocyte, Maurer’s cleft, and PVM membranes) or 0.1% Triton-X-100 (lyses all membranes). The pellet was separated from the supernatant (supe) and both fractions were incubated with PBS containing Proteinase K (PK) at 37 °C. Protease inhibitor cocktail and PMSF was added to all samples followed by reducing sample buffer before immunoblotting with anti-GFP (LSA3-GFP), anti-REX3 (exported control) or anti-aldolase (aldo; internal protein control for parasite membrane integrity) antibodies. LSA3-GFP is indicated in green while GFP core in the food vacuole is indicated. Representative data is from n=2-3 experiments.

Transfected parasites were validated by immunoblot using anti-GFP antibodies and live microscopy of the GFP signal. mLSA3-GFP had an apparent molecular weight of ∼29 kDa, consistent with proteolytic cleavage at the conserved leucine residue in the PEXEL motif by plasmepsin V and a smaller GFP core band derived from digestion of the reporter in the food vacuole, which confirmed it was secreted from the parasite (***Figure 3A, B***) (Boddey *et al*., 2009). In contrast, mLSA3 RLE>A-GFP had an apparent molecular weight of ∼39 kDa indicating cleavage of the N-terminus was inhibited by the point mutations and therefore cleavage was PEXEL-dependent, while the GFP core band was greatly diminished confirming the protein was not secreted correctly (***Figure 3B***) consistent with inhibited processing and retention inside the parasite (Boddey *et al*., 2010).

LSA3 is predicted to have an N-terminal hydrophobic region between residues 45 and 64 which likely targets the protein into the ER and secretory pathway. Putative cleavage after the hydrophobic sequence by signal peptidase would result in a fragment of 32 kDa yet this was not detected by immunoblot and SignalP 3.0 did not predict processing by signal peptidase; rather, it predicted an N-terminal signal anchor. To confirm that plasmepsin V was the protease responsible for N-terminal processing, parasites expressing mLSA3-GFP were incubated with either DMSO vehicle or the inhibitor WEHI-600 (Nguyen *et al*, 2018) and processing analysed by immunoblots. The band pattern observed for mLSA3-GFP in DMSO was consistent with N-terminal processing observed in the prior blot but incubation with WEHI-600 caused a larger precursor of mLSA3-GFP, of the same size as the PEXEL mutant mLSA3 RLE>A-GFP protein (***Figure 3B***). Therefore, plasmepsin V processed the N-terminus of mLSA3-GFP in a PEXEL-dependent manner during erythrocytic infection, providing direct evidence that this motif in LSA3 is a *bone fide* PEXEL, in agreement with its export to the erythrocyte.

Immunoblots suggested that inhibition of PEXEL processing of mLSA3 RLE>A prevented normal secretion from the parasite, as the quantity of the ‘GFP core’ degradation product was reduced. Micropscopy confirmed this result, with mLSA3 RLE>A-GFP being trapped within the parasite (***Figure 3C***). Past studies have shown that mutation of the PEXEL results in retention in the ER, suggesting that mLSA3 RLE>A-GFP may be trapped in this organelle (Boddey *et al*., 2009). Interestingly, while mLSA3-GFP containing the native PEXEL motif was cleaved and efficiently secreted out of the ER to the periphery (parasite membrane/PV/PVM), we did not detect export beyond the PVM for this reporter by IFA. This suggested that additional sequences downstream of the PEXEL that were not included in the minigene may be required for export, or that the addition of GFP influenced the final translocation of this protein across one or more membranes, which is unusual for *PEXEL* genes but not unprecedented, for example C-terminal mCherry blocked the export of LISP2 beyond the PVM of the infected hepatocyte (Orito *et al*., 2013) and LSA3-GFP below. Nonetheless, these findings demonstrate that mLSA3-GFP was N-terminally processed by plasmepsin V during asexual blood stage development and trafficked through the secretory pathway of the parasite, both in a PEXEL-dependent manner.

### Endogenous GFP tagging confirms LSA3 blood-stage expression but prevents export

While LSA3-C antibodies demonstrated expression of LSA3 in DGs (Morita *et al*., 2017) and in trophozoites (this study), we sought genetic confirmation of blood-stage expression of this liver-stage antigen. To this end, LSA3 was endogenously tagged with GFP at the C-terminus (***Figure 3D***) and visualised by live microscopy. Endogenous LSA3-GFP was expressed by blood-stage trophozoites and localized at the parasite periphery, suggestive of the PV / PVM. Interestingly, like the minigene mLSA3-GFP, no GFP signal was observed beyond the PVM (***Figure 3D***). This demonstrated by genetic epitope tagging that LSA3 is expressed during erythrocytic infection but that C-terminal fusion to GFP likely interfered with its export. Subcellular fractionation confirmed this to be the case, with LSA3-GFP present only in the pellet fraction after permeabilization of the erythrocyte membrane by EQT, and this was protected from PK digestion indicating it was not present in the erythrocyte (***Figure 3E***). Saponin permeabilization of the RBC membrane and PVM indicated further trafficking problems had occurred; firstly, a significant portion of LSA3-GFP was present in the saponin supernatant and could be degraded by PK, indicating it was soluble in the PV (***Figure 3E***). Secondly, the saponin pellet fraction of LSA3-GFP was resistant to PK, indicating the protein was located inside the parasite; based on the IFAs (***Figure 3D***), this was potentially at the plasma membrane with GFP facing internally (***Figure 3E***). Both results for LSA3-GFP were different to those observed after saponin fractionation of endogenous untagged LSA3 using LSA3-C antibodies, confirming that addition of GFP to the C-terminus affected both the trafficking and membrane association of LSA3, possibly by interfering with the C terminal hydrophobic sequence. We therefore did not study the LSA3-GFP parasite line any further. Interestingly, the GFP core of LSA3-GFP was consistently resistant to PK (***Figure 3E***), which agrees with the GFP fold also being resistant to proteases in the food vacuole, where the ‘GFP core’ species is formed following secretion from the parasite (Boddey *et al*., 2010; Boddey *et al*., 2009; Waller *et al*, 1999).

### LSA3-deficiency does not inhibit gametocytogenesis or transmission

To investigate the essentiality and function of LSA3 in *P. falciparum*, we assessed the development of Δ*LSA3* mutants in which the *LSA3* gene was disrupted (***Figure 1B***) across the lifecycle. NF54 and Δ*LSA3* clones D8 and E2 blood-stages were differentiated to gametocytes, and no significant differences were detected in the formation or quantity of stage V gametocytes across parasite lines (***Figure 4A***). Gametocytes were transmitted to *An. stephensi* mosquitoes by standard membrane feeding assays (SMFA) and midgut oocyst numbers were quantified 7 days postbloodmeal: again, no significant differences were observed for oocyst burden or infection prevalence between Δ*LSA3* clones and NF54 across independent experiments (***Figure 4B***). We next dissected the salivary glands of *An. stephensi* mosquitoes on day 17 postbloodmeal and quantified the number of sporozoites in separate cohorts; this revealed no significant differences in sporozoite yields between parasite lines (***Figure 4C***). Taken together, these results showed that LSA3-deficiency did not prevent development of mature gametocytes, transmission to or development of *P. falciparum* within mosquitoes or reduce the rate of sporozoite entry into the salivary glands under the conditions used.

**Figure 4.**
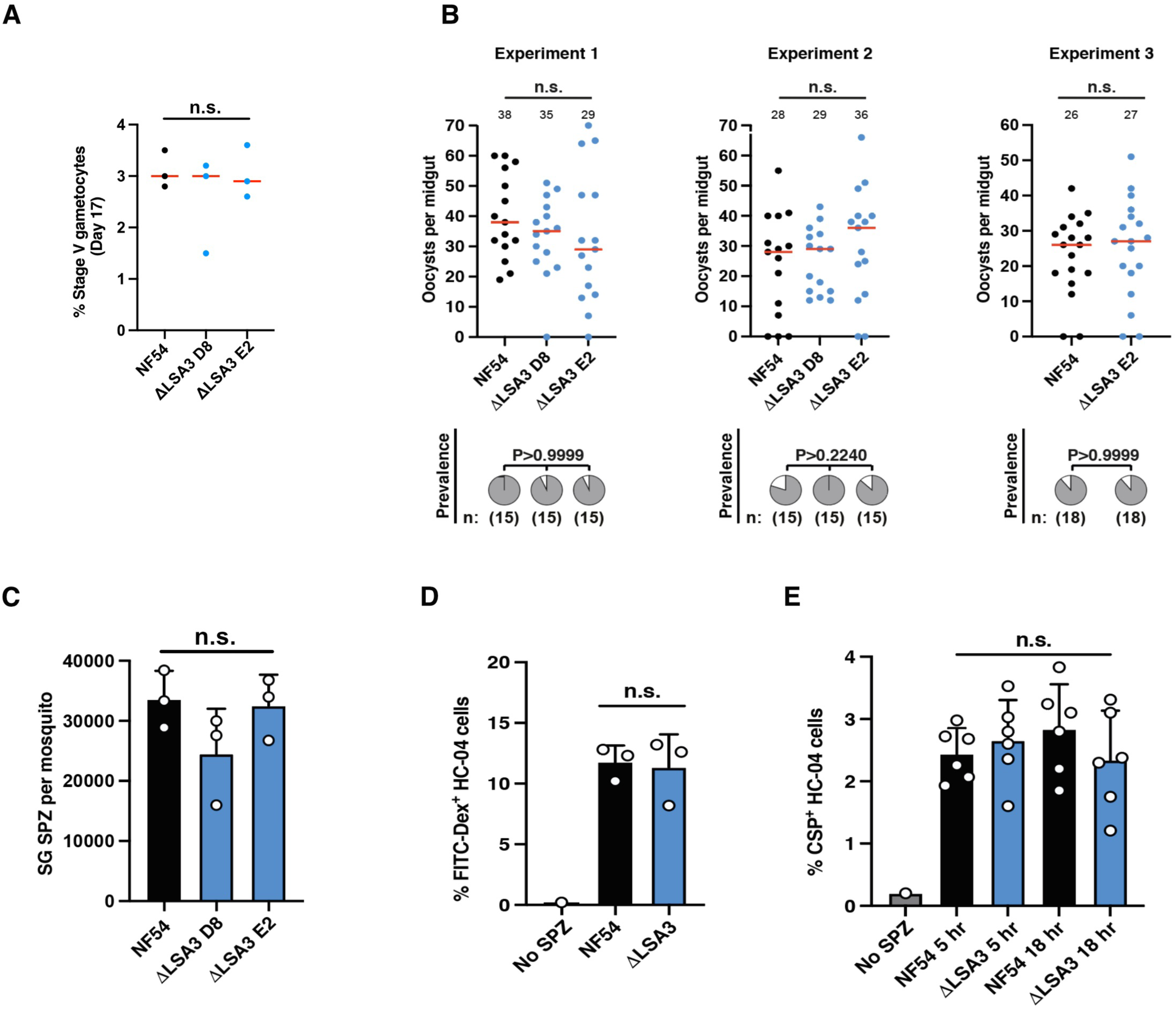
LSA3-deficiency does not inhibit gametocytogenesis or oocyst or sporozoite development nor sporozoite infectivity. **A.** Percentage of stage V gametocytes of NF54 parental and Δ*LSA3* clones D8 and E2. Red bars indicate the mean. Data was compared by ANOVA using the Kruskal Wallis test. **B**. Oocyst counts of NF54 and Δ*LSA3* clones D8 and E2 per mosquito midgut 7 days post bloodmeal. Data include the mean (number shown and red bar) from three independent transmission experiments. Data comparing Δ*LSA3* mutant clones to NF54 parent performed by ANOVA using the Kruskal Wallis test, except experiment 3 (Mann-Whitney test). Prevalence of mosquito infection is shown below each graph, comparing NF54 to Δ*LSA3* clones by chi-square analysis. **C.** Salivary gland sporozoite (SG SPZ) counts per mosquito at 17 days postbloodmeal. Data is mean compared Δ*LSA3* mutant clones to NF54 parents by ANOVA using the Kruskal Wallis test. **D.** Cell traversal enumerated as percentage of FITC-dextran^+^ HC-04 hepatocytes after incubation with freshy dissected NF54 and Δ*LSA3* clone E2 sporozoites from mosquito salivary glands. **E.** Percentage of HC-04 cells with CSP^+^ intracellular parasites at 5 and 18 hours of incubation with NF54 parent and Δ*LSA3* clone E2 sporozoites. Data was analyzed by ANOVA using the Kruskal Wallis test. All data in the figure is mean ± SEM from n=3 (A to D) or n=6 (E) independent experiments. In all panels, n.s., not significant (P>0.05). Data is from n=3 experiments.

### LSA3-deficiency does not perturb sporozoite traversal or infection of hepatocytes

Expression of LSA3 on the surface of *P. falciparum* sporozoites has been reported (Daubersies *et al*., 2000) suggesting it may be important in these forms. To investigate this, the kinetics of sporozoite traversal and infection of human hepatocytes were analysed. Firstly, cell traversal was investigated by incubating Δ*LSA3* clones or NF54 sporozoites with HC-04 human hepatocytes for 3 hours in the presence of the cell impermeant fluorescent molecule dextran-FITC. Only hepatocytes that have been traversed by sporozoites become permeable to dextran-FITC due to transient destabilization of the host cell membrane and these cells can be quantified by flow cytometry (Mota *et al*, 2001; Yang *et al*, 2017b). Both Δ*LSA3* clones and NF54 sporozoites were able to wound HC-04 hepatocytes leading to dextran-FITC uptake at similar rates (***Figure 4D***) indicating that LSA3 was not critical for sporozoite motility or their capacity to traverse human hepatocytes, which is a natural process of infection as sporozoites journey from the skin to blood vessels and eventually traverse hepatocytes before productively invading one in the liver.

We next investigated whether LSA3 was important for sporozoite infection of HC-04 hepatocytes using an assay that quantifies the number of intracellular parasites at different times of incubation for CSP-positive parasites by flow cytometry (Verzier *et al*, 2026). While CSP is initially located on the sporozoite surface, it is also present at the parasites periphery following human hepatocyte infection and during intracellular liver-stage development (McConville *et al*., 2024; Vaughan *et al*, 2012) and has been used to quantify productive invasion events on subsequent days (Angage *et al*, 2024; Kaushansky *et al*, 2015; Lopaticki *et al*, 2022; Verzier *et al*., 2026). We did not detect any significant difference in the number of CSP-positive Δ*LSA3* or NF54 parasites at 5 hours postinfection (hpi) or 18 hpi, indicating that LSA3 was not essential for *P. falciparum* sporozoite infection of HC-04 hepatocytes using this assay (***Figure 4E***).

### LSA3-C co-localizes with EXP1 and EXP2 in *P. falciparum* liver-stages

To better understand the localization of LSA3 at the liver stage, IFAs were performed on *P. falciparum* NF54 liver stages on days 5 and 6 post infection of humanized mice using the PVM markers EXP1 and EXP2. The mouse liver samples investigated were from a prior study in which humanized mice were singly infected with NF54 sporozoites (McConville *et al*., 2024). On day 5 postinfection, LSA3-C antibodies were detected around, and co-localizing with, the EXP1-labelled PVM (***Figure 5A***), and on day 6 with EXP2 at the PVM (***Figure 5B***). LSA3-C antibodies also localized in puncta beyond the circular shape of the EXP1-labelled PVM in a small number of infected hepatocytes; this was apparently LSA3 derived from egressing merozoites, or material released following PV rupture since LSA3-C was closely associated with the parasite DAPI signal (***Figure 5A***). These results confirm that LSA3-C antibodies localize at the PVM of *P. falciparum* exoerythrocytic forms. We did not detect LSA3-C beyond the PVM under the conditions tested. Whether LSA3 is exported in hepatocytes is uncertain: it may be considered exported if part of LSA3 reaches the host cell side of the PVM but confirming this will require follow up studies. As we employed a humanized mice co-infection strategy to assess the essentiality of Δ*LSA3* versus NF54, we could not yet perform IFAs on individually infected mice deficient for LSA3 to validate the specificity of LSA3-C or LSA3-T at the liver-stage.

**Figure 5.**
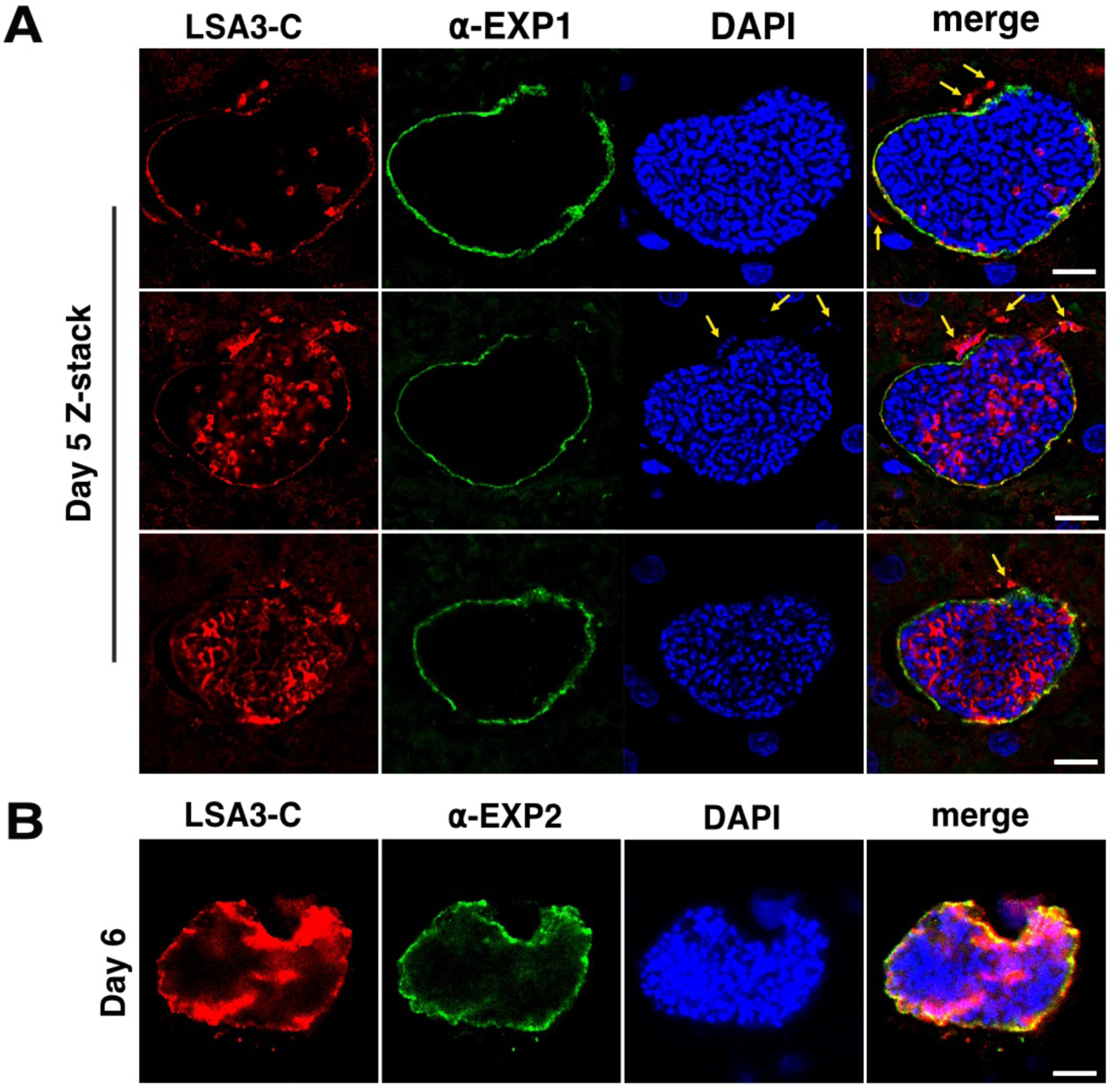
LSA3-C localizes to the PVM in *P. falciparum*-infected hepatocytes. **A.** Liver sections from humanized mice infected with *P. falciparum* NF54 sporozoites on day 5 post infection. Each panel shown is of the same liver-stage parasite at different cross-sections of the Z-stack. Sections were stained with anti-EXP1 antibodies that label the PVM and LSA3-C antibodies. LSA3-C localized predominantly to the PVM or with egressing merozoites beyond the PVM (yellow arrows). **B.** Liver section from humanized mice infected with *P. falciparum* NF54 sporozoites on day 6 post infection stained with anti-EXP2 antibodies that label the PVM and LSA3-C antibodies. For all images, parasite and host DNA was labelled with DAPI nuclear stain; scale bars, 10 µm. Representative images are from n=1 experiment from n=2 humanized mice.

### Genetic disruption of *LSA3* in *P. falciparum* attenuates liver-stage development

LSA3 has low sequence polymorphism across diverse geographical isolates and is a protective pre-erythrocytic antigen in chimpanzees and Aotus monkeys (Daubersies *et al*., 2000; Perlaza *et al*, 2003). To investigate its importance to parasite growth, Δ*LSA3* sporozoites were mixed in equal number with parental NF54 sporozoites and inoculated via a single intravenous (IV) injection into each of four SCID Alb-uPA humanized mice with chimeric human livers. At day 5 postinfection, livers were isolated from each mouse and half of the individual lobes were snap-frozen for sectioning and microscopy and the other half of each lobe was processed to recover genomic DNA for molecular quantification of parasite liver loads by real-time polymerase chain reaction (qRT-PCR) (Foquet *et al*, 2013) using oligonucleotide pairs that amplified DNA specific to either Δ*LSA3* or NF54 from the same livers. This experimental approach allows the fitness of mutant parasites to be directly compared to parental controls in the same mice (Lopaticki *et al*, 2017; McConville *et al*., 2024; Yang *et al*, 2017a). The day 5 endpoint was chosen as this coincided with the peak of PVM localization reported previously (Daubersies *et al*., 2000) and before parasites egress from the liver on day 7. This analysis revealed a 2.3-fold reduction in the liver load for Δ*LSA3* compared to NF54 on day 5 post infection (P=0.0110) (***Figure 6A***). When represented as a percentage of total parasite liver load, where equal fitness of each co-infected parasite line corresponds to 50%, Δ*LSA3* averaged a 40% reduction in fitness compared to NF54 (P=0.0105) (***Figure 6B***). The four humanized mice had similar levels of human chimerism of approximately 20% human:80% mouse hepatocytes, as determined by qRT-PCR (***Figure 6C***). Together, these results show that genetic disruption of *LSA3* in *P. falciparum* caused a moderate and significant loss of fitness in each of four humanized mice, indicating that this PEXEL-containing protein is important for normal intrahepatic development of the human malaria parasite.

**Figure 6.**
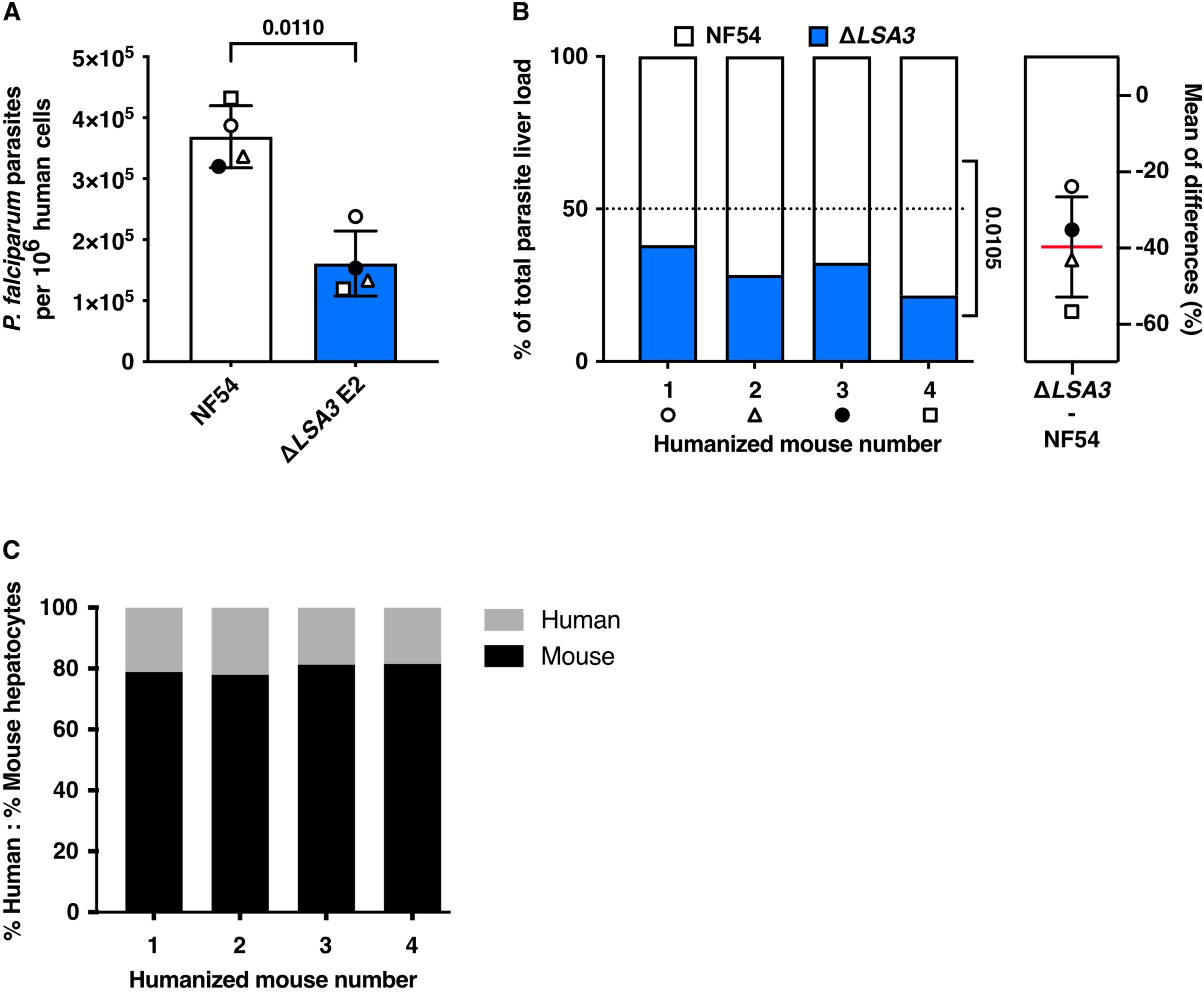
LSA3 is important for *P. falciparum* liver-stage development in humanized mice. **A.** Quantification of *P. falciparum* genomes per million human hepatocytes on day 5 postinfection from four co-infected humanized mice with chimeric human livers, each receiving 5×10^5^ Δ*LSA3* clone E2 and 5×10^5^ parental NF54 sporozoites mixed for a total of 1×10^6^ sporozoites via a single IV injection per mouse, by qRT-PCR. Values from each sample were normalized to a series of pretested standard curves of each amplicon and are shown as mean ± SD, statistically analysed by paired *t*-test. Each symbol corresponds to the same coinfected mouse. **B.** Percent of total parasite liver load on day 5 postinfection in each humanized mouse co-infected with equal numbers of Δ*LSA3* E2 and parental NF54 sporozoites from (A), shows reduced fitness of LSA3-deficient parasites (i.e. less than 50%). Percent differences of the mean in each mouse are shown on the right ± SD. Each symbol corresponds to the same coinfected mouse. Means were compared by paired *t*-test. **C.** Human chimerism in each humanized mouse (21%, 22%, 19% and 18%) quantified by qRT-PCR using oligonucleotides specific to human and mouse PTGER2. Data is from n=1 experiment from n=4 humanized mice.

## Discussion

Historically, LSA3 was considered a protein expressed only in pre-erythrocytic stages (Daubersies *et al*., 2000), but more recent work identified LSA3 in merozoite DGs during RBC invasion (Morita *et al*., 2025; Morita *et al*., 2017). Here, endogenous GFP tagging confirmed LSA3 expression in blood stages and motivated biochemical and functional studies across the parasite lifecycle, prompted by a previously uncharacterized PEXEL motif (Maier *et al*., 2008; McConville *et al*., 2024; Morita *et al*., 2017). LSA3 is processed by plasmepsin V (inhibited by a PEXEL-mimetic inhibitor) and is associated with the PVM, while a fraction is exported into the infected erythrocyte as soluble aggregates distinct from Maurer’s clefts. Its biochemical profile resembles that of the exported chaperone HSP70-x, suggesting possible association with J-dots or other chaperone complexes (Hanssen *et al*, 2008; Kulzer *et al*, 2012; Kulzer *et al*., 2010; Petersen *et al*, 2016; Taraschi *et al*, 2003). AlphaFold predicts an LSA3 domain with similarity to the DnaK substrate binding domain (Jumper *et al*., 2021), consistent with a chaperone-like role. Limitations in fixation and antibody specificity so far precluded definitive colocalization with HSP70-x; identifying interacting partners will help clarify LSA3’s role in blood-stages and may inform its liver-stage function.

LSA3’s DG localization aligns with discharge during merozoite invasion, when DG contents contribute to PV and PTEX assembly and early export (Morita *et al*., 2017; Riglar *et al*, 2013). Given its DG association and recent blood-stage vaccine interest, LSA3 may act during or immediately after invasion (Morita *et al*., 2025; Morita *et al*., 2017). Trafficking of LSA3 reporter proteins required PEXEL processing by plasmepsin V, but C-terminal fluorescent fusions disrupted correct targeting, and endogenous tagging showed predominate retention at the parasite plasma membrane and a soluble PV pool, consistent with the sensitivity of pre-erythrocytic trafficking to bulky tags (Orito *et al*., 2013; Singer & Frischknecht, 2021).

Understanding of *P. falciparum* liver-stage biology remains limited, yet exported proteins expressed at this stage are plausible antigenic and functional candidates. LSA3 antibodies localized mainly to the PVM in infected hepatocytes with little evidence of export beyond the membrane. Recent reports suggest PEXEL processing can direct proteins to rhoptries or the PV in blood stages (Fierro *et al*., 2024; Freville *et al*., 2024) and the related apicomplexan *Toxoplasma,* uses an analogous TEXEL/ASP5 pathway to target to the PV or host cell compartments (Bougdour *et al*, 2014; Coffey *et al*, 2015; Hammoudi *et al*, 2015). Whether a domain of LSA3 is exported into the hepatocyte lumen from the PVM remains unresolved.

In this study, genetic disruption of *LSA3* reduced liver-stage fitness by 40%, indicating an important role for LSA3 during hepatocyte infection. This makes LSA3 the second PEXEL protein required for normal *P. falciparum* liver-stage development, after LISP2, which causes arrest later when genetically disrupted (Zanghi *et al*., 2025). The differential localization between blood and liver stages suggests stage-specific routing of PEXEL proteins: in blood stages PEXEL processing commonly precedes export to the host cell cytosol whereas in liver stages PVM targeting is more frequent (Fougere *et al*, 2016; Ingmundson *et al*, 2012; McConville *et al*., 2024). Hence, the export pathway may influence PVM remodelling and host-pathogen interactions during liver-stage development (Liehl *et al*., 2014; Marques-da-Silva *et al*., 2025; Prado *et al*., 2015).

LSA3 is immunogenic at the liver stage (Daubersies *et al*., 2000), and more recently at the blood-stage (Morita *et al*., 2017), and its sequence conservation across geographical isolates suggests functional constraint. Structural prediction revealed similarity to a DnaK-like substrate binding domain, and a large unstructured repeat region, with different geographical isolates showing variation in the number of repeats (Prieur & Druilhe, 2009). Unstructured repeats are common features of other *Plasmodium* proteins (Orito *et al*., 2013) and disordered regions feature in exported proteins in Apicomplexa (Bougdour *et al*, 2013; Dumaine *et al*, 2021; Huang *et al*, 2022; Rodrigues *et al*, 2025; Rosenberg & Sibley, 2021).

In summary, LSA3 is a component of the *P. falciparum* exportome, required for efficient liver-stage development and capable of eliciting protective pre-erythrocytic immunity. Defining its precise functional roles at the PVM and in the host erythrocyte will help inform antigen discovery while identification of additional exported proteins trafficked to these locations may facilitate vaccine design and reveal novel drug targets.

## Materials and methods

### Parasite maintenance

*P. falciparum* NF54 (kindly provided by the Walter Reid Army Institute of Research) asexual stages were cultured in human O-positive erythrocytes (Melbourne Red Cross) at 4% haematocrit in RPMI-HEPES supplemented with 0.2% NaHCO_3_, 7% heat-inactivated human serum (Melbourne Red Cross), and 3% AlbuMAX (ThermoFisher Scientific). Parasites were maintained at 37 °C in 94 % N_2_, 5 % CO_2_ and 1 % O_2_. Gametocytes were cultured in RPMI-HEPES supplemented with 0.2 % NaHCO_3_, and 10 % heat-inactivated human serum (Melbourne Red Cross). Gametocytes for transmission to mosquitoes were generated using the “crash” method (Saliba & Jacobs-Lorena, 2013) using daily media changes.

### Generation of transgenic *P. falciparum*

The *LSA3* knockout clones Δ*LSA3* (E2) and (D8) were generated by amplification of the 5’ and 3’ flanks of the *LSA3* gene (PF3D7_0220000). Oligonucleotides used were: forward primer 5′-AAACCGCGGATGTAGATAAGAAATTGAATAAAC-3′ and reverse primer 5′-TTGAACTAGTATTTCTTCAACACTTGGAGC-3′ (5′ flank); forward primer 5′-TTAGAATTCAAAAATATGGAAGAGGAGTTAATGAAG-3′ and reverse primer 5′-TTACCTAGGTCTTCATCTATATCTTCATCTATATC-3′ (3′ flank). The flanks were ligated on either side of the human DHFR cassette in pCC1 (see *Figure 1*). Highly synchronous *P. falciparum* NF54 schizonts were enriched by magnet-activated cell sorting (MACS) magnetic separation columns (Miltenyi biotec) and incubated with E64 (Sigma) until parasitophorous vacuole (PV) enclosed merozoite structures (PEMS) could be visualized (Boyle *et al*, 2010). Purified *LSA3* knockout plasmid DNA (100 μg, Life Technologies) was transfected into the schizonts with an Amaxa Basic Parasite Nucleofector Kit 2 (Lonza). Parasites that had correctly integrated the knockout construct were selected by cycling on and off 4nM WR99210 and stable transfectants were cloned out by limiting dilution (Voss *et al*, 2006) and confirmed by diagnostic PCR.

### Mosquito infection and analysis of parasite development

Seven-day old female *Anopheles stephensi* mosquitoes were fed on asynchronous gametocytes diluted to 0.6% stage V gametocytemia, via water jacketed glass membrane feeders as previously described (McConville *et al*., 2024). Mosquitoes were sugar starved for 48 hours following feeding to enrich for blood fed mosquitoes. Surviving mosquitoes were provided with di-ionised water via paper/cotton wicks and sugar cubes. Oocyst numbers were obtained from midguts dissected from cold-anesthetized and ethanol killed mosquitoes 7 days post feed and stained with 0.1% mercurochrome. Salivary glands were dissected from mosquitoes (day 16 to 20 post bloodmeal), crushed using pestle and then glass wool filtered to obtain sporozoites that were quantified using a hemocytometer and used in subsequent assays.

### Immunofluorescence microscopy assay (IFA)

Asexual stages were fixed in 4% paraformaldehyde (PFA) PBS with 0.0075% glutaraldehyde, permeabilised with 0.01% triton X 100 or fixed and permeabilised with ice cold 90% Acetone 10% methanol and probed with primary antibodies; mouse anti-GFP (Roche; Cat # 11814460001; RRID:AB_390913; 1:1000), mouse anti-MAHRP2 (1:1000) (Pachlatko *et al*., 2010), mouse anti-EXP2 (1:500) (de Koning-Ward *et al*., 2009), mouse anti-REX3 (1:1000) (Spielmann *et al*., 2006), and rabbit anti-LSA3-C (1:1000) (Morita *et al*., 2017) or LSA3-T (1:1000) (Morita *et al*., 2025), in 3% normal donkey serum (NDS/ PBS). Secondary antibodies were goat anti-rabbit Alexa 594 and anti-mouse or anti-rat Alexa 488 (Thermo Fisher; Cat # A-11012, A-11029, A-11006; RRID:AB_2534079, RRID:AB_2534088, RRID:AB_2534074; 1:1000). For liver-stage: chimeric human livers were perfused with 1x PBS and fixed in 4% (vol/vol) PFA PBS overnight before being exchanged for sucrose and snap frozen in OCT (Invitrogen). Sections of 8 mm were cut on the cryostat apparatus HMM550. Samples were permeabilised using ice cold 10% methanol 90% acetone and blocked using 3% BSA PBS. Sections infected with NF54 parasites were then incubated with primary antibodies: rabbit anti-LSA3-C (1:1000), mouse anti-PfCSP (2A10; 1:2000) (Nicholas *et al*, 2023), rat anti-EXP1 (McConville *et al*., 2024) (1:1000), mouse anti-EXP2 (1:500), diluted in 3% BSA PBS. Secondary antibodies were the same goat anti-rabbit Alexa 594 and anti-mouse or rat Alexa 488 (1:1000). Samples were incubated with DAPI (4’,6’-diamidino-2-phenylindole) at 1 mg/ml in PBS to visualize DNA and mounted in Prolong Gold mounting media (Invitrogen). For live imaging, blood stage culture was incubated with DAPI for 2 min, washed once with PBS and 4 uL blood pallet dropped onto cover slips and inverted. Images were acquired immediately. All images were acquired on the Zeiss LSM 980 microscope.

### Subcellular fractionation, proteinase K treatment and immunoblots

Late-stage parasites at desired stage purified by Percoll gradient were resuspend in 40 ul PBS + 5 ul EQTII (0.5mg/ml) per 5 ul of pellet and incubate at RT for 6 min. The supernatant was collected; pellet washed in PBS and resuspend in the same volume as the supe before suspension of fractions in 4x SDS-PAGE loading buffer and stored at -20°C. For Saponin, Percoll-purified parasites were resuspended in 45 uL 0.03% saponin, incubated at RT for 3 min. Supernatant was collected, the pellet washed in PBS and resuspend in the same volume as the supe before suspension of fractions in 4x SDS-PAGE loading buffer and stored at -20°C. For Triton X-100, Percoll-purified parasites were resuspend in 45 uL 0.1% TX-100, incubated at RT for 3 min. Supernatant was collected, the pellet washed in PBS and resuspend in the same volume as the supe before suspension of fractions in 4x SDS-PAGE loading buffer and stored at -20°C (Gabriela *et al*., 2022). Proteinase K solution (20 ug/mL; Worthington) was added to samples, incubated at 37°C for 20 mins and then samples resuspended in 4x SDS-PAGE loading buffer. Proteins were separated through 4-12% Bis-Tris gels and transferred to nitrocellulose before blocking in 5% milk PBS-T and probing with mouse anti-GFP (Roche; Cat # 11814460001; RRID:AB_390913; 1:1000), mouse anti-REX3 (1:1000), rabbit anti-aldolase (Baum *et al*, 2006) (1:4000) followed by anti-mouse (1:1000) or anti-rabbit (1:4000) horse radish peroxidase-conjugated secondary IgG antibodies (Cell Signalling Technology; Cat # 7076S, 7074S; RRID:AB_330924, RRID:AB_2099233) and viewed by enhanced chemiluminescence (Amersham) (Jennison *et al*, 2019).

### LSA3 sequence analysis and generation of the LSA3 minigene

The analysis of LSA3 from *P. falciparum* 3D7 was performed by inputting the accession Pf3D7_0220000 into <u>PlasmoDB</u> or <u>AlphaFold</u>. The first 112 amino acids of the LSA3 coding sequence were synthesised by Integrated DNA Technologies containing either the endogenous PEXEL sequence or one where the key residues, RLE, were mutated to alanines. These were then ligated into the pKiwi003 (Ashdown *et al*., 2020) using restriction sites NheI and PstI, for integration into the *P. falciparum* 3D7 *P230* locus. The resulting plasmid was transfected into PEMs schizonts in conjunction with Cas9-U6.2-hDHFR_P230p (50 ug of each) following standard procedures (Boyle *et al*., 2010) and correct integration was selected for following addition of 4 nM WR99210 for 10 days.

### Plasmepsin V cleavage inhibition assay

*P. falciparum* parasites at 24-34h old trophozoites were purified from uninfected erythrocytes Vario Macs magnet column (Miltenyi Biotech) separation and treated with 20 uM of the plasmepsin V inhibitor WEHI-600 (Nguyen *et al*., 2018) or DMSO vehicle for 3 h at 37°C in culture gas. Parasites were treated with 0.1% saponin to liberate contaminating hemoglobin and pellets solubilized in 4x Laemmli sample buffer before protein separation via SDS-PAGE. Proteins were separated through 4-12% Bis-Tris gels and transferred to nitrocellulose before blocking in 5% milk PBS-T and probing with mouse anti-GFP (Roche; Cat # 11814460001; RRID:AB_390913; 1:1000), followed by anti-mouse (1:1000) or anti-rabbit (1:4000) horse radish peroxidase-conjugated secondary IgG antibodies (Cell Signalling Technology; Cat # 7076S, 7074S; RRID:AB_330924, RRID:AB_2099233) and viewed by enhanced chemiluminescence (Amersham) (Sleebs *et al*, 2014).

### Hepatocyte culturing

HC-04 hepatocytes (Sattabongkot *et al*, 2006) were maintained on Iscove’s Modified Dulbecco’s medium (IMDM), supplemented with 5% heat-inactivated fetal bovine serum (FBS) at 37°C in 5% CO_2_. Cells were split 1:6 every 2-3 days once they reached 90% confluency.

### Cell traversal assays

Cell traversal was measured using a cell wounding assay (Lopaticki *et al*., 2022). HC-04 hepatocytes (1×10^5^) were seeded onto the bottom of a 96-well plate using IMDM (Life Technologies, 11360-070). Sporozoites dissected from mosquito salivary glands into Schneider’s insect medium were enumerated and placed into IMDM containing 10% human serum before addition at an MOI 0.3 to hepatocyte monolayers for 2.5 hr for cell traversal to occur in the presence of 1 mg ml-1 FITC-labelled dextran (10,000 MW, Sigma Aldrich). Cells were washed and trypsinized to obtain a single cell suspension for FACS quantification of dextran-positive and -negative cells. For each condition, triplicate samples of 10,000 cells were counted by flow cytometry in each of the three independent experiments.

### Hepatocyte invasion assays

HC-04 hepatocyte invasion assays were performed as previously described (Lopaticki *et al*., 2022). Briefly, HC-04 hepatocytes (1×10^5^) were seeded onto the bottom of 96-well plates using IMDM (Life Technologies, 11360-070). Sporozoites dissected from mosquito salivary glands into Schneider’s insect medium were quantified and replaced in IMDM containing 10% human serum before addition at MOI 0.3 to hepatocytes for either 5 or 18 hr before being washed and trypsinized to obtain a single cell suspension and transferred to a 96 well round bottom plate. Cells were fixed and permeabilised (BD bioscience) and stained with Alexa fluor 647 conjugated mouse anti-PfCSP antibodies in 3% BSA permeabilizing solution. Cells were then washed in PBS and analysed by flow cytometry to quantify CSP^+^ HC-04 cells. For each condition, triplicate samples of 10,000 cells were counted in each of the three independent experiments.

### Humanized mice production and processing

uPA^+/+^ SCID mice (University of Alberta) were housed in a virus and antigen free facility supported by the Health Sciences Laboratory Animal Service at the University of Alberta and cared for in accordance with the Canadian Council on Animal Care guidelines. All protocols involving mice were reviewed and approved by the university of Alberta Health Sciences Laboratory Animal Service Animal Welfare committee and the Walter and Eliza Hall Institute of Medical Research Animal Ethics Committee. uPA+/+-SCID mice at 5–14 days old received 10^6^ human hepatocytes (cryopreserved human hepatocytes were obtained from BioreclamationIVT—Baltimore MD) by intrasplenic injection and engraftment was confirmed 8 weeks following transplantation by analysis of serum human albumin (Mercer *et al*, 2001; Yang *et al*., 2017b). For humanized mice co-infections: 5 × 10^5^ Δ*LSA3* clone E2 and 5 × 10^5^ parental NF54 sporozoites (1×10^6^ total) were injected IV into each of 4 mice. Livers were isolated 5 days postinfection from CO_2_-ethanized mice and individual lobes were cut as described (Lopaticki *et al*., 2017), pooled, and emulsified into a single cell suspension and flash frozen in liquid nitrogen for subsequent genomic DNA (gDNA) extraction.

### Measuring exoerythrocytic development in humanized mice

To quantify parasite load in the chimeric livers, gDNA was isolated from the single cell liver suspensions and Taqman probe-based qPCRs were performed as previously described (Alcoser *et al*, 2011; Lopaticki *et al*., 2017; Yang *et al*., 2017b). To specifically differentiate NF54 from Δ*LSA3* clone E2 genomes from the same mouse samples, oligonucleotides were designed as below. Human and mouse genomes were quantified using oligonucleotides specific for prostaglandin E receptor 2 (PTGER2) from each species, as described previously (Yang *et al*., 2017b). All probes were labelled 5′ with the fluorophore 6-carboxy-fluorescein (FAM) and contain a double-quencher that includes an internal ZENTM quencher and a 3′ Iowa Black® quencher from IDT. The following probes were used:

LSA3WT-F: TCCGCTAATTTCTACTGTTTCCTCT

LSA3WT-R: GAACAGGCAGAAGAAGAGAGC

LSA3WT: FAM/TGCAGAGAA/ZEN/TTTAGAGAA-3IABkFQ

hDHFR_F 5′-ACCTAATAGAAATA TATCAGGATCC-3′

hDHFR_R 5′-GTTTAAGATGGCCTGG GTGA-3′

*ΔLSA3* (DHFR): FAM/CCGCTCAGG/ZEN/AACGAATTTAGATA-3IABkFQ

Pf18s-F: GTAATTGGAATGATAGGAATTTACAGGT

Pf18s-R: TCAACTACGAACGTTTTAACTGCAAC

18S: FAM/TGCCAGCAG/ZEN/CCGCGGTA-3IABkFQ

hPTGER2: FAM/TGCTGCTTC/ZEN/TCATTGTCTCG/3IABkFQ

mPTGER2: FAM/CCTGCTGCT/ZEN/TATCGTGGCTG/3IABkFQ

Standard curves were prepared by titration from a defined number of DNA copies for each amplicon of *P. falciparum* LSA3WT, Δ*LSA3*, 18S, and human PTGER2 and mouse PTGER2 controls. PCRs were performed on a Roche LC80 using LightCycler 480 Probe Master (Roche) (Lopaticki *et al*., 2017).

### Statistics

Parasite conditions were compared between groups using an ANOVA Kruskal-Wallis test with Dunn’s correction, or Mann Whitney test where indicated, whilst mosquito infection prevalence was compared using the chi-square test. Statistical analyses comparing Δ*LSA3* clone E2 to NF54 controls in co-infected mice were performed using the paired t-test and comparisons of individual mutants to control were performed using the Mann-Whitney test. Analyses were performed using Graphpad Prism 9 for macOS. P<0.05 was considered statistically significant.

### Ethics statement

Experimental protocols involving humanized mice were conducted in accordance with the recommendations in the National Statement on Ethical Conduct in Animal Research of the National Health and Medical Research Council and were reviewed and approved by the University of Alberta Health Sciences Animal Welfare Committee and the Walter and Eliza Hall Institute of Medical Research Animal Ethics Committee. Experimental protocols involving the HC-04 human hepatocyte cell line were reviewed and approved by the Walter and Eliza Hall Institute of Medical Research Biosafety Committee.

## Acknowledgements

We thank the Melbourne Red Cross for human erythrocytes, the US Naval Medical Research Centre for the human HC-04 hepatocyte cell line, Joao Aguiar for EXP1 antibodies, Paul Gilson for EXP2 antibodies, Eizo Takashima for LSA3-C and LSA3-T antibodies, Matt Dixon for REX3 antibodies, Leanne Tilley for EQTII and MAHRP2 antibodies, Fidel Zavala for PfCSP 2A10 antibodies, and Jake Baum for pKIWI construct. We acknowledge Julie Healer and Melissa Hobbs for technical assistance in the insectary.

## Additional information

### Author contributions

Experiments were conceptualized by R.M. and J.A.B. and performed by R.M., R.W.J.S., and M.T.O. Methodology was performed by R.M., M.T.O., N.K., and J.A.B. Data was interpreted by R.M., A.F.C., and J.A.B. R.M. and J.A.B. drafted, and all authors contributed to preparing and editing, this manuscript.

### Data and materials availability

All data and materials generated in this study are available from the corresponding author upon request.

---

**Figure S1.**
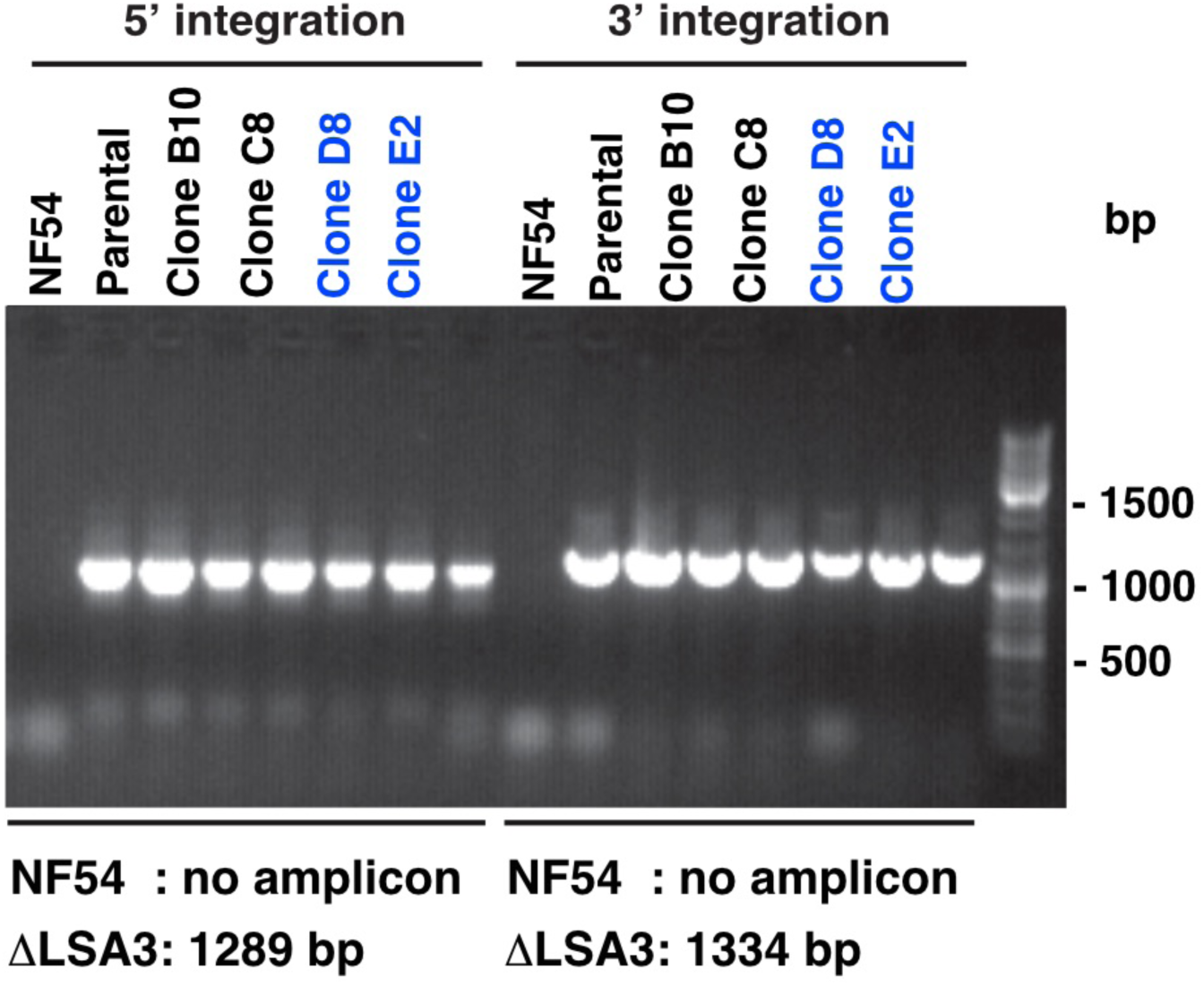
Genotyping *P. falciparum* NF54 and Δ*LSA3* clones. Diagnostic PCRs (MO545/aw560) confirmed correct double cross-over integration of the Δ*LSA3* knockout pCC1 construct in *P. falciparum* NF54 clones. Mutant clones D8 and E2 used in this study are shown in blue and molecular weight sizes in base-pairs (bp) are shown on the right.

**Figure S2.**
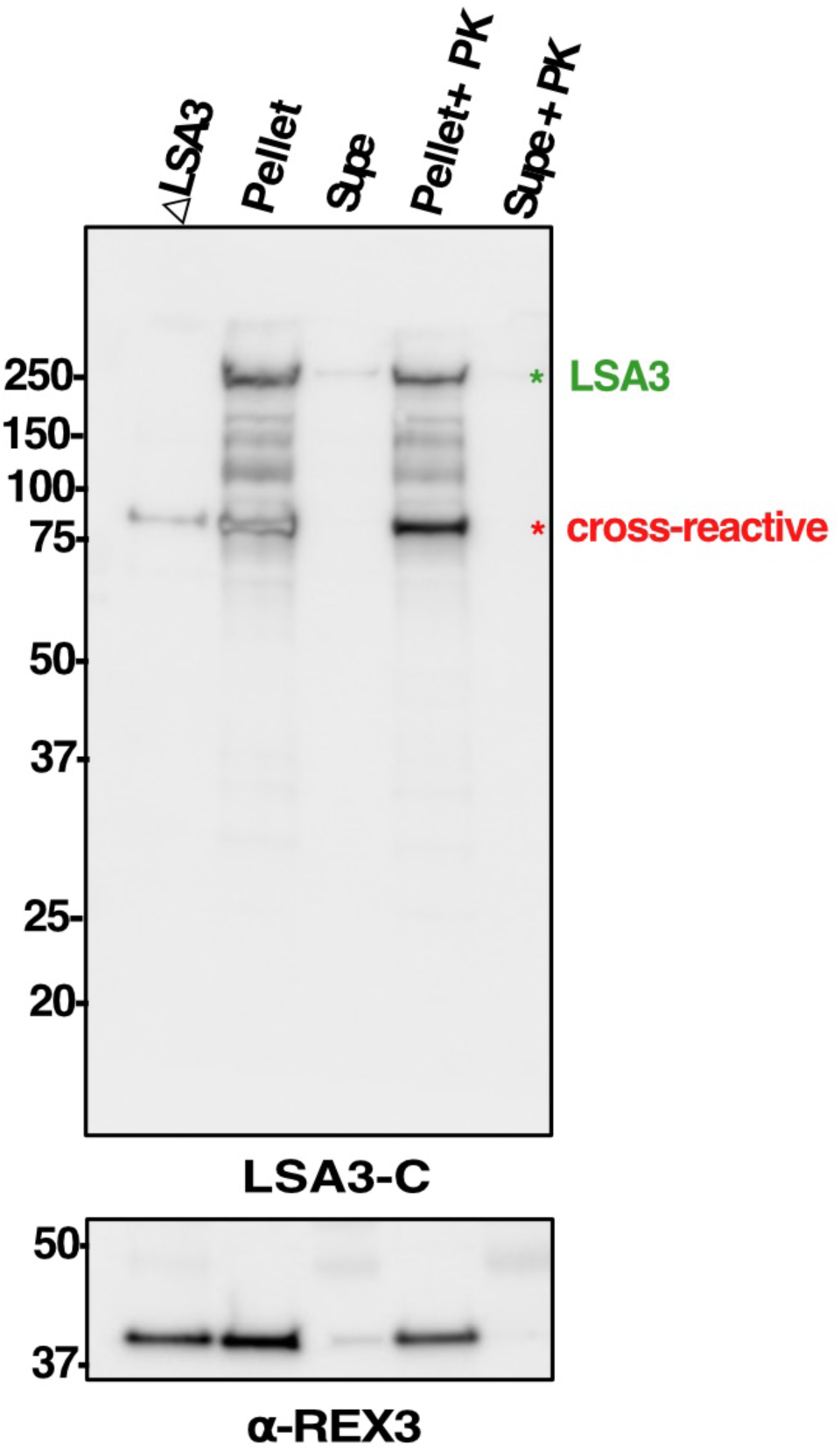
The LSA3-C binding domain is not localized on the infected erythrocyte surface. Erythrocytes infected with Δ*LSA3* and NF54 parental parasites were enriched by Percoll centrifugation, with the former being included as a specificity control for the LSA3-C antibody. NF54-infected erythrocytes were incubated with PBS to maintain cells as intact. The pellet was then separated from the supernatant (supe) fraction by centrifugation, and both were incubated at 37 °C with Proteinase K (PK) in PBS. Protease inhibitor cocktail and PMSF were added to all samples followed by reducing sample buffer to stop the reactions. Samples were separated by SDS-PAGE and probed with LSA3-C or anti-REX3 (exported protein control) antibodies. The C-terminal portion of LSA3 recognized by LSA3-C antibodies was not present on the erythrocyte surface as it remained resistant to Proteinase K, like the exported REX3 control as expected. The cross-reactive internal protein recognized by LSA3-C is visible in all lanes.

